# Dynamical models reveal distance to criticality in ageing brain dynamics

**DOI:** 10.1101/2025.09.26.678711

**Authors:** Vivek Sharma, Bas J. N. M. Drost, Paul H. E. Tiesinga

**Author notes:** These authors contributed equally to this work (Vivek Sharma, Bas J. N. M. Drost). Neurophysics, Donders Institute for Brain, Cognition and Behaviour, Radboud University, Nijmegen, Netherlands (Paul H. E. Tiesinga).

## Abstract

Understanding how the brain changes with age remains a central question in neuroscience. Here, we combine magnetoencephalography (MEG) recordings from young and older adults with a whole-brain dynamical model to explore how brain dynamics evolve across the lifespan. Using a network of coupled Stuart-Landau oscillators constrained by empirical structural connectivity, we systematically vary three model parameters to identify the settings that best reproduce alpha-band features observed in MEG data. Our findings reveal age-related shifts in these model parameters: older individuals exhibit stronger global coupling and more positive values of the bifurcation parameter, consistent with a transition to a supercritical regime. These results align with prior work suggesting altered excitation-inhibition balance in ageing and indicate a systematic reconfiguration of whole-brain dynamics. By situating empirical observations within a dynamical systems framework, this study provides a principled approach for quantifying the brain’s distance to criticality and lays the groundwork for future clinical applications.

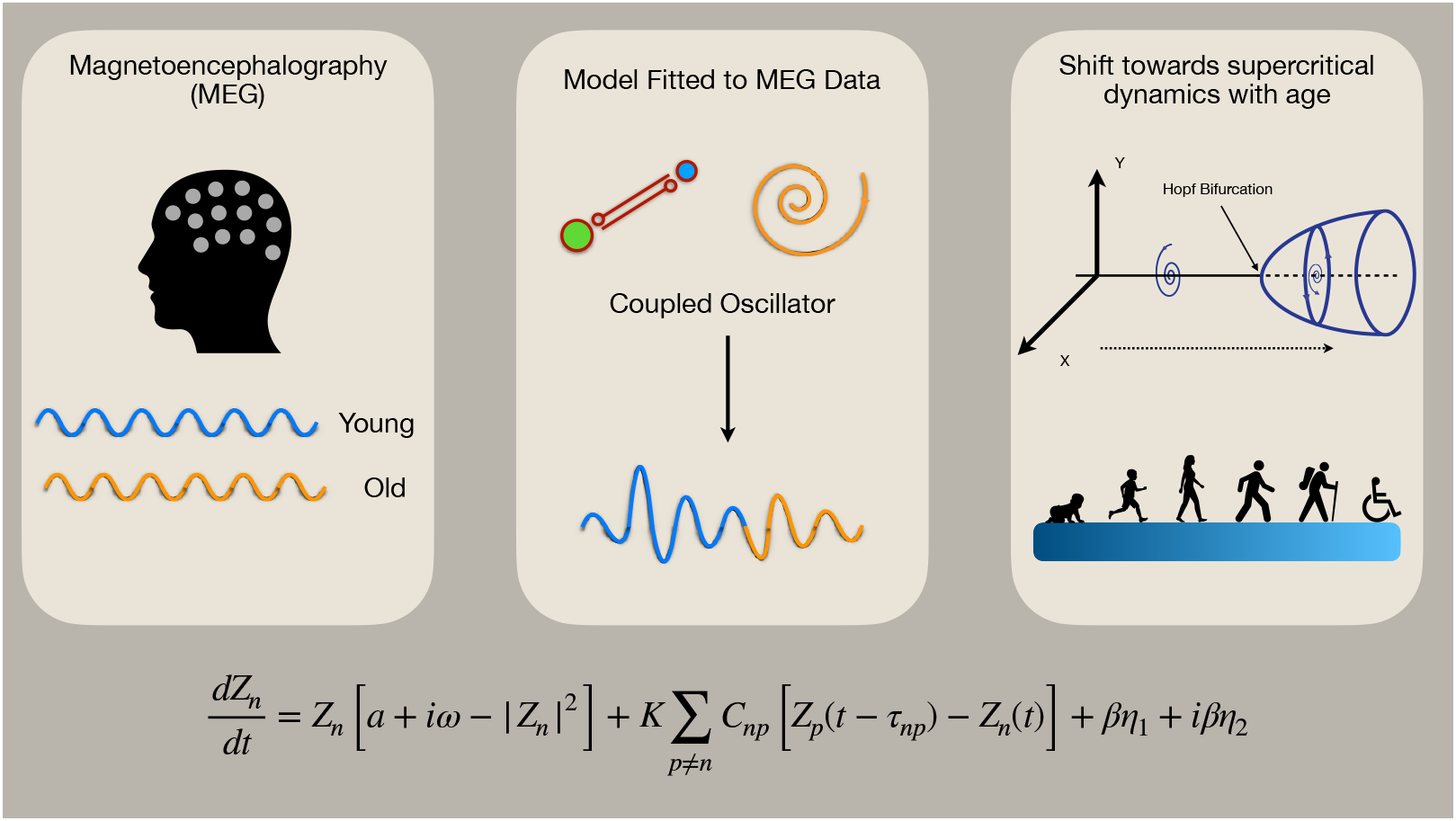

## Introduction

Understanding how brain dynamics evolve throughout the lifespan remains a central challenge in neuroscience. Recent studies employing magnetoencephalography (MEG) and electroencephalography (EEG) have consistently demonstrated significant age-related alterations in neural oscillations, network connectivity, and the excitation-inhibition (E/I) balance, particularly within the alpha frequency band (8–13 Hz) [1, 2]. Young adults typically display robust alpha-band activity, higher peak alpha frequencies, and strong alpha-band coupling within and between resting-state networks, which contrasts with the diminished alpha power, shift in peak alpha, and weaker functional connectivity often observed in older adults [1–5].

Alpha oscillations and functional connectivity exhibit characteristic developmental trajectories. Studies report an inverted U-shaped trajectory, with alpha-band power and peak frequency reaching their maxima during early adulthood before progressively declining with age [6]. Accompanying these oscillatory changes, older adults demonstrate increases in beta power and altered transient network dynamics. Specifically, the occurrence of early visual states decreases with age, while the occurrence of fronto-temporo-parietal and higher-order visual states increases. This age-related reorganisation parallels earlier findings of altered temporal stability of dynamic networks across development [7], and in older adults, it is more strongly associated with poorer fluid intelligence (the capacity to reason and solve novel problems), suggesting reduced neural efficiency rather than functional compensation [5].

Computational and whole-brain models provide mechanistic insights into these empirical observations. Models utilising coupled oscillators constrained by population-averaged structural connectivity suggest that the brain compensates for structural decline by increased conduction delays through enhanced inter-areal coupling, thereby preserving neural synchrony [1, 8]. Furthermore, computational analyses reveal age-associated shifts in the excitation-inhibition balance, where older brains move towards increased excitation, perhaps explaining both neural noise and cognitive performance decrements observed in older age groups [6, 9].

Crucially, the concept of criticality, the state in which neural systems are optimally positioned between order and disorder, offers a theoretical framework to interpret these age-dependent changes. A system near criticality demonstrates maximal responsiveness and adaptability, which is essential for healthy cognitive function [10]. However, ageing may shift brain dynamics beyond this critical state into a supercritical regime characterised by sustained oscillatory activity and reduced flexibility, driven by altered excitation-inhibition balance [1, 6].

Building on this foundation, the present study aims to elucidate how ageing influences the brain’s proximity to criticality using a whole-brain modeling framework constrained by structural connectivity. By systematically varying key parameters such as the value of the bifurcation parameter, global coupling strength, and mean conduction delays, we seek to closely replicate empirical MEG data from young and older adults. Our goal is to capture the functional consequences of ageing on brain-wide neural activity and provide a quantitative framework for applying distance-to-criticality metrics within clinical neuroscience contexts.

## Methods and Materials

### 0.1 Participants

Data for this study were sourced from the Cam-CAN repository, a population-based, multi-modal, cross-sectional investigation spanning adult lifespan [11, 12]. The study protocol was approved by the Research Ethics Committee of Cambridgeshire 2, with informed written consent from all participants. Initially, 2681 participants underwent home-based interviews and assessments covering neuropsychological functions, sensory capabilities (vision, hearing), balance, and response speed. Exclusion criteria included impaired vision, hearing loss greater than 35 dB at 1000 Hz, substance abuse history, psychiatric disorders (e.g., bipolar disorder, schizophrenia), neurological conditions (e.g., epilepsy, stroke, traumatic brain injury), or Mini-Mental State Examination scores below 25. Subsequently, 700 eligible participants advanced to Stage 2; from these, magnetoencephalography (MEG) data were available for 650 individuals.

### 0.2 Data Acquisition

The MEG recordings analysed in this study were obtained from the publicly available Cam-CAN dataset (http://www.mrc-cbu.cam.ac.uk/datasets/camcan/). Data acquisition was performed using an Elekta Neuromag system (Helsinki) equipped with 306 sensors, comprising 102 magnetometers and 204 planar gradiometers. Recordings took place in a magnetically shielded room under dim lighting, with signals sampled at 1000 Hz and filtered online with a 0.03 Hz high-pass. Head position was continuously tracked using head-position indicator (HPI) coils, while horizontal and vertical EOG electrodes captured eye blinks and movements. Each participant underwent at least 8 minutes and 40 seconds of eyes-closed resting-state MEG recording.

### 0.3 Data Pre-processing

The Cam-CAN project provided preprocessed MEG data, which had already undergone a standardised cleaning pipeline. This included temporal signal space separation (tSSS) to suppress artifacts arising from HPI coils, environmental noise, and continuous head motion. A MaxFilter procedure was further applied to attenuate line noise (50 Hz notch filter) and to interpolate or reconstruct noisy channels. Full details of the acquisition and preprocessing protocols are available in Cam-CAN et al. (2014) and Taylor et al. (2017) [11, 12].

### 0.4 Data Analysis

Participants were grouped into younger (18–34 years) and older adults (66–88 years), each comprising 50 individuals [13]. Source localisation utilised the Colin27 anatomical template, segmented with FreeSurfer [14], and sources were reconstructed with standardised Low-Resolution Brain Electromagnetic Tomography (sLORETA) in MNE-Python [15, 16]. The forward model was computed via the Boundary Element Method (BEM), and source-space definitions were established following Desikan-Killiany cortical parcellation [17]. Source time series were subsequently filtered to the alpha frequency range (8–13 Hz).

### 0.5 Structural Connectivity

Structural connectivity (SC) matrices were constructed from diffusion-weighted MRI data obtained from the 1000BRAINS Cohort, which includes 261 healthy participants aged 18–86 years. Connectivity was defined using the Desikan-Killiany parcellation scheme [20, 21]. The population-averaged SC matrix was used to constrain the model dynamics, providing a normative baseline that enhances reliability while reducing subject-level noise [22, 23].

### 0.6 Functional Connectivity

Functional connectivity (FC) was quantified using amplitude envelope correlation (AEC) [24, 25]. AEC assesses the correlation between amplitude envelopes derived from bandpass-filtered MEG signals, providing a robust measure of neural coupling. FC matrices were computed per participant, then averaged within age groups to form empirical references for model fitting.

### 0.7 Simulations

Whole-brain simulations employed the Stuart-Landau oscillator model, chosen for its representational simplicity and capacity to capture neural dynamics near criticality [23]. Oscillatory dynamics at each brain region were modelled using the Stuart-Landau Eq (1):

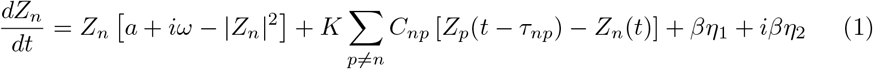

Here, *Z*_*n*_ represents the complex state of node *n*, with intrinsic frequency (*ω*), bifurcation parameter (*a*), global coupling (*K*), and connectivity (*C*). Time delays (*τ*_*np*_) were calculated based on fibre lengths and conduction speed (*v*). Gaussian noise (*η*) was included to simulate neural variability. Model parameters (*K*, mean delay, and *a*) underwent a systematic grid search (9,240 parameter combinations, each simulated 30 times) to comprehensively explore neural dynamics.

### 0.8 Model Fitting and Analysis

Simulation accuracy was evaluated by comparing simulated FC to empirical FC using Pearson correlation. Parameter sets yielding the highest correlations within alpha-band constraints (7–13 Hz) were identified. Subsequently, specific alpha sub-bands (10–13 Hz for young, 7–8.5 Hz for old) were isolated to differentiate age-related dynamics clearly. The final analyses were based on the top 200 fitting scores per age group, ensuring robust comparison between groups. Figure 1 illustrates the overall model fitting pipeline, from structural connectomes and MEG data to functional connectivity, fitting scores, and power spectral density analyses.

**Fig 1.**
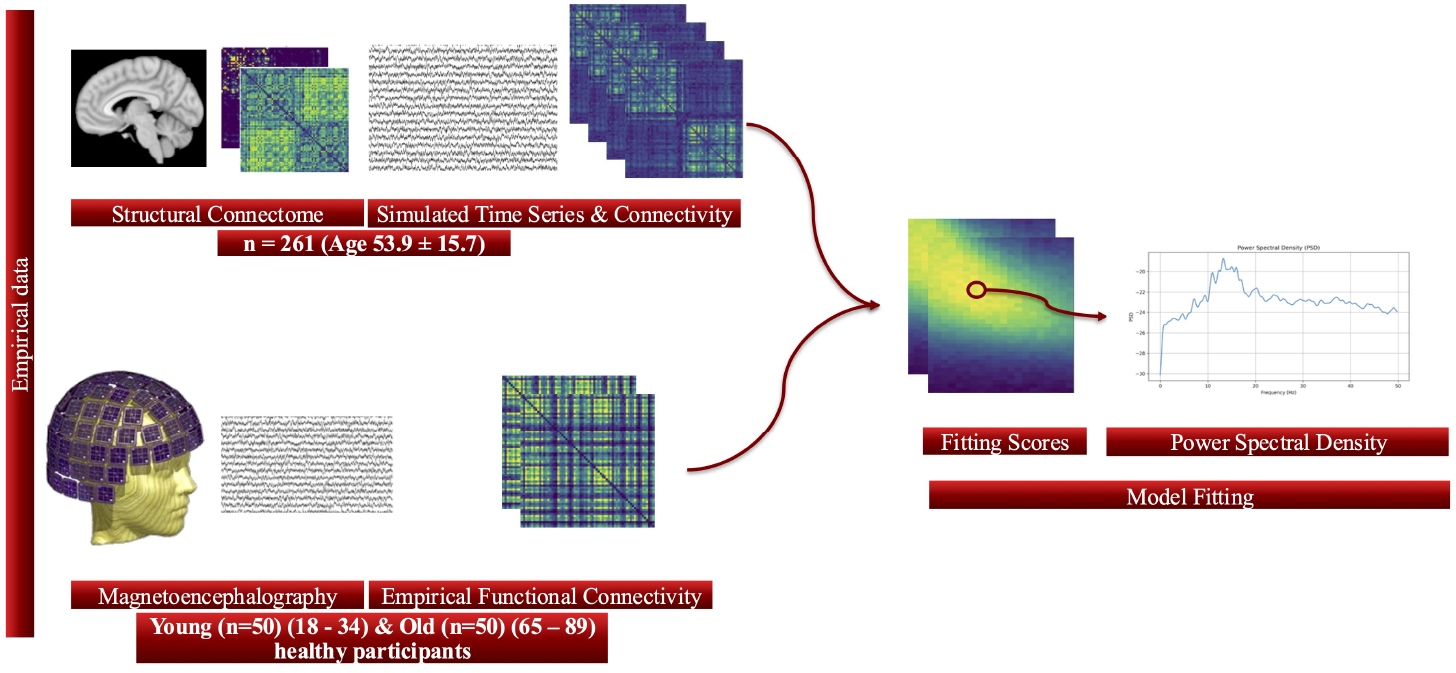
Schematic overview of the model fitting. Structural connectomes (n = 261, mean age 53.9 ± 15.7 years) were used to simulate large-scale brain dynamics, generating time series and functional connectivity. Magnetoencephalography (MEG) data from healthy young (n = 50, 18–34 years) and old (n = 50, 65–89 years) participants provided empirical functional connectivity and spectral properties. Model parameters were optimised through fitting simulated to empirical connectivity, and evaluated using power spectral density and fitting scores, enabling the characterisation of critical brain dynamics across age groups. Some panel elements were adapted from [18] and [19].

### 0.9 Statistical Analysis

All group comparisons were performed using the non-parametric Mann–Whitney U test, as the parameter distributions violated normality assumptions. For each model parameter and outcome metric, we report the p-values associated with group differences. To control for multiple comparisons across the eight primary variables (e.g., model fit score, alpha-band metrics, and model parameters), we applied the Benjamini–Hochberg procedure for false discovery rate (FDR) correction. Violin plots in Fig. 2 display the distributions of each metric by age group, along with raw p-values and FDR-corrected significance levels. All statistical analyses were conducted in Python using SciPy and statsmodels libraries.

**Fig 2.**
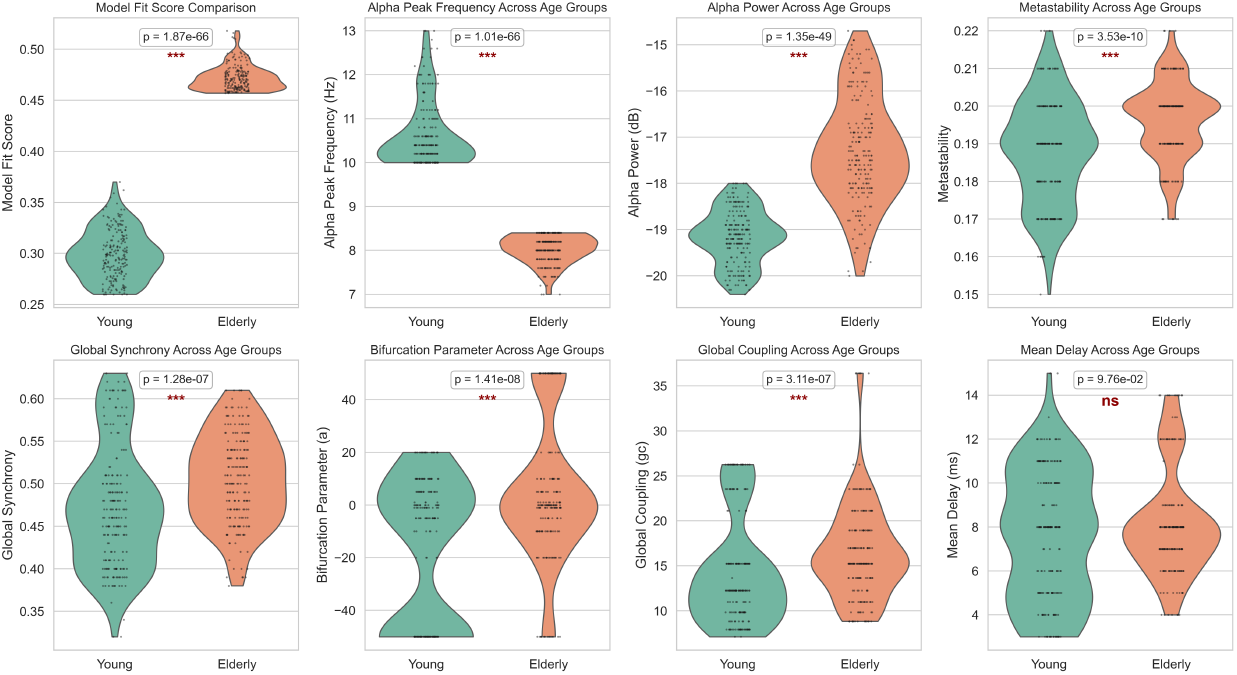
Group-level comparison of model fit scores and model parameters between younger and older participants. Violin plots show the distribution of each variable within the young (green) and elderly (orange) groups across eight metrics: model fit score, alpha peak frequency, alpha power, metastability, global synchrony, bifurcation parameter (*a*), global coupling (*K*), and mean delay (*md*). Each violin includes a swarm plot overlay representing individual subjects. Statistical significance of between-group differences was assessed using the Mann–Whitney U test; p-values and significance levels (*, **, ***) are annotated above each panel. Notably, the elderly group exhibited significantly higher model fit scores, global coupling, alpha power, and a shift in the bifurcation parameter toward the supercritical regime, indicating altered large-scale brain dynamics with ageing.

## Results

### 0.10 Age-Dependent Shift in Critical Brain Dynamics

A pronounced shift in dynamical regime was observed between younger and older participants, most clearly reflected in the distribution of the bifurcation parameter *a* (Fig. 2). In the younger group, the majority of best-fitting solutions were in the subcritical regime, with 62.5% of individuals exhibiting *a <* 0 and with a group mean of *a* = −18.0. This places younger adults predominantly in the subcritical regime, where dynamics are fluctuation-dominated, noise-sensitive, and poised between quiescence and oscillation. In contrast, the elderly group displayed a marked shift toward supercritical dynamics, with a mean *a* = 4.45 and only 43% of individuals retaining *a <* 0. This transition suggests a fundamental reconfiguration of brain dynamics with age, consistent with a departure from criticality and the emergence of self-sustained oscillatory behaviour.

### 0.11 Differences in Model Fit and Biophysical Parameters

Model fit scores were significantly higher in the elderly group (mean = 0.47 ± 0.00087) compared to the young group (mean = 0.30 ± 0.0017), indicating that the model more accurately captured the empirical functional connectivity of older individuals. Global coupling strength (*K*) was also elevated in the elderly (mean = 16.4 ± 0.37 vs. 14.25 ± 0.41 in young), potentially reflecting increased long-range integration or compensatory network reorganisation.

Alpha power was higher in elderly participants (−17.29 dB) relative to the young group (−19.16 dB), consistent with previous findings of enhanced oscillatory amplitude in ageing. The mean delay parameter (*md*) was slightly longer in the elderly (8.20 ± 0.17 ms vs. 7.61 ± 0.21 ms), although variance was greater in the younger cohort (SD = 2.93 ms vs. 2.44 ms), suggesting more heterogeneous conduction dynamics in youth.

Other network-level metrics, such as metastability and global synchrony, showed only modest group differences, with a trend toward more stable, less variable dynamics in the elderly brain.

### 0.12 Emergent Frequency Dynamics and Critical Coupling

Across all simulations, we observed a nonlinear interaction between global coupling (*K*), mean delay (*md*), and the emergent alpha peak frequency (Fig. 3). Specifically, increasing delay was associated with progressively slower oscillations, while a sharp frequency transition occurred near a critical coupling threshold (*K* ≈ 5.71) when *md >* 10 ms. Beyond this threshold, the alpha frequency dropped abruptly, indicating a bifurcation-like behaviour in frequency dynamics.

**Fig 3.**
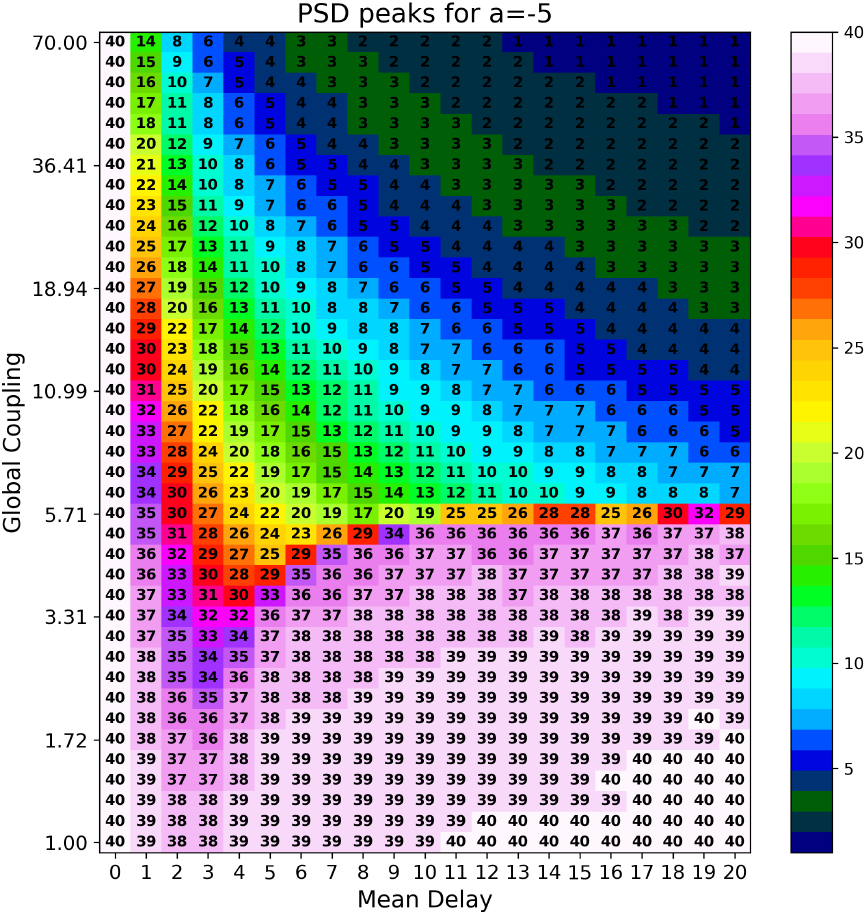
PSD peak frequency as a function of mean delay and global coupling (*a* = −5). Each grid cell shows the dominant frequency (Hz) obtained from simulations for the corresponding parameter set. Increasing mean delay (*md*) is associated with slower peak frequencies, while a sharp transition occurs at a critical coupling strength (*K* ≈ 5.71) when *md >* 10 ms, beyond which the system’s dominant frequency drops abruptly.

Additional simulations revealed that this critical *K* threshold varies with the bifurcation parameter *a*, with more excitable local dynamics (higher *a*) requiring greater coupling to induce global synchrony. These results demonstrate that the brain’s operating frequency is not fixed, but emerges from a delicate balance between local excitability and global communication delays, a balance that shifts systematically with age.

## Discussion

The present study provides a robust investigation into how ageing affects brain dynamics, particularly through the lens of criticality, excitation-inhibition (E/I) balance, and alpha-band oscillations. By combining empirical magnetoencephalography (MEG) data and computational whole-brain modeling using Stuart-Landau oscillators, we delineate clear age-related shifts in neural dynamics, consistent with recent findings from both neuroimaging and computational modeling literature [1, 3, 4, 6].

A key finding is the shift in bifurcation parameter (*a*) observed with age may reflect an underlying change in excitation-inhibition balance, where more positive values suggest a transition toward sustained, supercritical oscillations. Biophysically, this aligns with increased cortical excitation or reduced GABAergic inhibition—consistent with less negative spectral exponents seen in EEG studies of older adults [9]. Meanwhile, global coupling (*K*) can be interpreted as a proxy for large-scale synchronisation and compensatory plasticity, potentially reflecting recruitment of additional networks to maintain integration despite declining structural integrity [1]. Mean delay (*md*), though more stable, may be associated with demyelination and reduced axonal conduction velocity, as reported in age-related white matter degeneration [1].

The critical coupling threshold identified in our analyses further highlights the complex interplay between global network connectivity and local neural excitability. The distinct transition observed at a coupling strength around *K* = 5.71 underlines the sensitivity of the neural system to specific parameter conditions, consistent with theoretical predictions of phase transitions in dynamical systems [23]. This critical point is also characterised by increased variability in peak frequency, a hallmark of critical slowing down typically observed near bifurcation transitions [23].

Structural connectivity plays a pivotal role in shaping brain dynamics. Our model, based on population-averaged SC matrices, robustly captured age-related differences; however, future work using individualised SC data may refine these distinctions further. Such personalisation could help uncover subtle inter-individual variations in the ageing process and better guide the development of dynamical biomarkers. [22, 23].

These findings resonate with multiple recent empirical and computational studies [1, 3–5, 26]. Ageing consistently associates with decreased alpha-band power, reduced alpha coupling, and lower peak frequencies, paralleled by increased excitation and heightened beta-band activity. Such dynamics appear to reflect both neural degradation and compensatory reorganisation, evident through enhanced coupling mechanisms designed to preserve neural synchrony despite declining structural connectivity and increased axonal delays [1, 9].

While our work frames criticality in terms of proximity to bifurcations, ageing also alters statistical (power-law) signatures of brain activity. EEG studies report that older adults show larger increases in DFA (detrended fluctuation analysis) scaling exponents (i.e., stronger long-range temporal correlations) during motor tasks relative to young adults [27]. Resting-state fMRI shows a reduction of the power-law trend in the global signal with age, alongside a shift of power from lower to higher frequencies (“temporal dedifferentiation”) [28]. Complementing these, MEG analyses within a quasicriticality framework on the Cam-CAN lifespan cohort reveal age-related correlations in critical-like metrics, linking ageing, connectivity changes, and increased dynamical fluctuations [29]. Aligning these results suggests that ageing impacts both the dynamical (bifurcation-centred) and statistical (power-law) facets of criticality in the brain.

Nevertheless, our study has some inherent limitations. The use of population-averaged SC rather than individualised connectomes may obscure subject-specific differences in network structure and function. While the averaged approach reduces noise and highlights generalised network characteristics, future work would benefit from individual-level SC data to capture more granular distinctions. Furthermore, the cross-sectional design limits our ability to infer causal relationships or dynamic developmental trajectories with confidence.

Despite these limitations, the results presented here significantly advance our understanding of age-related changes in brain dynamics. Our approach illustrates how whole-brain computational models, combined with high-quality empirical neuroimaging data, can provide meaningful insights into fundamental neural mechanisms altered by ageing. This framework may offer powerful biomarkers for early detection of neurodegenerative disorders and inform targeted interventions aimed at restoring optimal E/I balance and network criticality. Future studies should explore longitudinal data and expand computational methods to further elucidate how neural dynamics evolve throughout the lifespan, potentially guiding novel therapeutic strategies in clinical neuroscience. These findings open avenues for clinical applications. For instance, distance-to-criticality metrics may serve as early biomarkers of cognitive decline or neurodegeneration, particularly in conditions such as Alzheimer’s disease. Furthermore, understanding individual proximity to criticality could inform personalised neurostimulation protocols (e.g., tACS/tDCS) aimed at restoring optimal network responsiveness.

## Acknowledgments

eBRAIN-Health has received funding from the European Union’s Horizon Europe research and innovation programme under grant agreement No 101058516.

